# Graphic insights into the network basis for heat-gated TRPV1

**DOI:** 10.1101/2022.01.02.474701

**Authors:** Guangyu Wang

## Abstract

Non-covalent interactions in bio-macromolecules are individually weak but collectively important. How they take a concerted action in a complex biochemical reaction network to realize their thermal stability and activity is still challenging to study. Here graph theory was used to investigate how the temperature-dependent non-covalent interactions as identified in the structures of the thermo-gated capsaicin receptor TRPV1 could form a systemic fluidic grid-like mesh network with topological grids to maintain the 3D structure of TRPV1 and to govern heat-sensing. The results showed that the heat-evoked melting of the biggest grid may initiate a specific temperature threshold for both channel activation and inactivation. Meanwhile, a thermostable balance between an open state and a putatively inactivated state from the same pre-open closed state may account for heat efficacy and the use-dependent desensitization, which were further stabilized by smaller grids. Altogether, both the biggest and smaller grids may be necessary for the temperature sensitivity. Therefore, this grid thermodynamic model may be broadly significant for the structural thermostability and the functional thermoactivity of biological macromolecules.

**Highlights:**

- Thermo-driven cyclization or decyclization of a DNA hairpin and a network grid in protein was thermodynamically comparable
- The temperature thresholds of TRPV1 for channel activation and inactivation were comparable theoretically and experimentally
- The thermo-sensitivities of TRPV1 for channel activation and inactivation were comparable theoretically and experimentally
- The grid-based systemic thermal instability of TRPV1 was useful to identify different gating states
- The release of the phosphatidylinositol lipid from the vanilloid site was required for the heat-evoked activation of TRPV1

## 1. INTRODUCTION

Temperature is very important in everyday life. Transient receptor potential (TRP) vanilloid-1 (TRPV1) is well-known as a noxious heat sensor in a primary afferent neuron of the dorsal root and the trigeminal ganglia. Its activation allows sodium or calcium to be permeable to a channel pore. With an activation threshold (∼42 °C) above a normal human body temperature 37 °C and a temperature coefficient Q_10_ (the ratio of rates or open probabilities (P_o_) of an ion channel measured 10 °C apart) higher than 20, this channel can work as a bio-thermometer in a thermosensory system to monitor a noxious heat signal above 42 °C, protecting human or animal bodies from heat damage [1-4]. In addition to the physical stimuli, capsaicin can also activate TRPV1, producing hot sensation. As cold can decrease the capsaicin-evoked activity of TRPV1 [5], the same gating pathway seems to be shared by both stimuli. On the other hand, residues sensitive to both physical and chemical stimuli are different in TRPV1, suggesting their distinct gating pathways [6-9].

TRPV1 is a homotetramer with each monomer consisting of six transmembrane helices. The first four segments (S1–S4) form a voltage-sensor-like domain (VSLD) while the last two (S5 and S6), together with the intervening loop and the pore helix and the extracellular cap-like pore turret between S5 and S6, are folded as a pore domain. The pore domain is surrounded by the VSLD in a domain-swapped arrangement. The ion permeation pathway centers in the pore domain. A hydrophobic constriction around I697 forms a lower S6 gate while the segment from G643 to D646 (^643^GMGD^646^) between S5 and S6 (S5-P-S6) in rat TRPV1 (rTRPV1) acts as a short dynamic ion selectivity filter. The S4–S5 linker, which is a short amphipathic helix and almost parallel to the cytoplasmic membrane, couples the VSLD with the pore domain for channel gating. Thus, the overall structure of the TRPV1 channel resembles that of the voltage-gated ion channel [10-11]. However, TRPV1 has a conserved TRP domain, which runs parallel to the inner leaflet of the membrane by virtue of a sharp bend after S6, not only linking the S6 gate crossing bundle but also interacting with the S4–S5 linker and the pre-S1 domain in favor of allosteric coupling between different gating domains. In addition, the characteristic ankyrin repeats within the cytoplasmic N terminus can tether cytoplasmic N- and C-terminal domains for channel assembly [11-14]. Although the structural basis for the chemical stimuli such as capsaicin has been clear [6, 8, 15-17], little is known about the structural factors or motifs to control the heat-evoked activation and use-dependent desensitization of capsaicin receptor TRPV1 above 42 °C and related temperature sensitivity [1, 8, 18-20].

Since 2013 a DNA hairpin thermal sensor with a limited poly-A loop length has been created to monitor a change in a biological microenvironment. When temperature increases above a threshold in a range from 34 °C to 73 °C, the hydrogen bonds between the bases in the DNA duplex are broken so that the DNA hairpin melts with a characteristic sigmoid plot [21]. Similarly, TRPV1 can be activated above a specific temperature threshold 42 °C and also with a featured sigmoid plot at elevated temperatures [18]. Because a grid in a grid-like non-covalent interaction mesh network in protein is topologically similar to a DNA hairpin in general term and the dissociation of this network grid by heat is analogous to DNA hairpin melting, it is fitting to ask if the thermosensitive TRPV1 channel exploits such topological network grids to keep the 3D structure of TRPV1 for heat sensing and subsequent desensitization.

Recently, graph theory, together with the temperature-dependent structural and functional data of DNA hairpin thermosensors, has been successfully used to establish a grid thermodynamic model with two powerful equations to shed light on the network basis for the structural thermostability or the functional thermoactivity of a biological macromolecule, class I aldolase B [22]. One was to calculate the melting temperature threshold (T_m_) of the biggest grid as the threshold (T_th_) for the changes in the tertiary or secondary structure or the specific catalytic activity; the other was to evaluate the systematic thermal instability (T_i_) as an important energetic reference of the structure-activity relationship. Hence, these two equations could be employed here to define temperature-dependent gating states and to identify the structural motifs or factors for heat-sensing and the use-dependent desensitization.

On the other hand, it has been reported that the opening rate or the open probability (P_o_) of TRPV1 is increased at raised temperatures [19, 23]. Thus, if the temperature threshold for TRPV1 opening is determined by a rate-limiting single step to disrupt a non-covalent interaction in the biggest grid along the gating pathway, TRPV1 opening would be enthalpy-driven (ΔH<0) because the dissociation rate (*k_d_*) of enthalpy-driven non-covalent interaction in a biophysical network is factually accelerated at elevated temperatures but *k_d_* of entropy-driven crosslinks is negligibly affected or even slowed under these conditions [24]. In this case, when a thermo-gated TRPV1 opens from a closed state within 10 °C, the functional thermo-sensitivity or the temperature coefficient (Q_10_) should be governed by the enthalpy-driven changes of the whole grids in the non-covalently interacting mesh networks. In this regard, once the structural thermo-sensitivity (Ω_10_) was similarly and approximately defined as a heat-evoked change of the total chemical potential of all the grids upon a change in the total enthalpy included in non-covalent interactions along the same gating pathway of a single polypeptide chain between open and closed or inactivated states within 10 °C apart, Ω_10_ should be comparable to Q_10_.

Taken together, these three lines of computational and comparative results favored three proposals. First, the melting temperature threshold (T_m_) of the biggest grid may be responsible for the specific temperature threshold (T_th_) for TRPV1 activation and inactivation; Second, a thermostable balance between an open state and a putatively inactivated state from the same pre-open closed state may determine the heat efficacy and the use-dependent desensitization with a reasonable energetic sequence; Third, the temperature coefficient Q_10_ of TRPV1 may be controlled by the grid-based structural thermo-sensitivity Ω_10_.

## 2. METHODS

### 2.1 Data mining resources

In this computational study, graph theory was used to analyze the topological grids in a systemic fluidic grid-like non-covalently interacting mesh networks of rTRPV1 below and above the temperature thresholds as the molecular thermometer basis for noxious heat detection. Those networks were based on the cryo-EM structural data of closed rTRPV1 in MSP2N2 with PI-bound at 4 **°**C (PDB ID, 7LP9, model resolution = 2.63 Å) and 48 **°**C (PDB ID, 7LPC, model resolution = 3.07 Å), capsaicin-bound at 25 **°**C (PDB ID, 7LPB, model resolution = 3.54 Å) and 48 **°**C for 10 s (EMD ID, 23477, model resolution = 3.70 Å) and 30 s (EMD 23478; PDB ID, 7LPD, model resolution = 3.55 Å), and open rTRPV1 with capsaicin-bound at 48 **°**C (PDB ID, 7LPE, model resolution = 3.72 Å) [14].

### 2.2 Standards to filter non-covalent interactions

UCSF Chimera was used to assess the stereo- or regioselective inter-domain diagonal and intra-domain lateral non-covalent interactions such as salt-bridges, lone pair/CH/cation-π interactions and H-bonds between amino acid side chains along the gating pathway from D388 in the pre-S1 domain to K710 in the TRP domain of rTRPV1. Their consistent standard definition as described previously was employed as a filter to secure that results could be reproduced with a high sensitivity [22]. Further in this study, when a distance between a cation and an aromatic center was within 6.0 Å, significant cation-π interactions were also considered. In contrast, the lone pair-π interaction distance between a lone electron pair and an aromatic ring was within 3-3.7 Å.

### 2.3 Preparation of topological grid maps by using graph theory

After non-covalent interactions were filtered along the gating pathway of rTRPV1 from D388 in the pre-S1 domain to K710 in the TRP domain, a grid was defined according to the graphical method as described previously [22]. All the temperature-dependent grids were geometrically mapped along the gating pathway in the well-established closed and open states and in the putatively desensitized state. The map had a black line to represent the primary amino acid sequence from D388 to K710. The colorful arrow nodes represented amino acid side chains or atoms of a bound lipid for diffferent non-covalent interactions along the gating pathway of rTRPV1. The colorful edges between nodes represented distinct non-covalent interactions in a grid-like biochemical reaction mesh network. The same kind of non-covalent interactions was marked in the same color. A topological grid was formed between two nodes *i* and *j* (n*_i_* and n*_j_*), if and only if there were a path from n*_i_* to n*_j_* and a return path from n*_j_* to n*_i_*. The grid size (S) was definded as the minimal number of the total side chains of residues in protein or atoms in the bound lipid that did not involve any non-covalent interaction in a grid. For a given grid-like biochemical reaction mesh network, the grid size between n*_i_* and n*_j_* was the sum of the shortest return path distance from node *j* to *i* because the direct shortest path distance from node *i* to *j* was zero when a non-covalent interaction existed between them. The Floyd-Warshall algorithm was used to identify the shortest return path between two nodes *i* and *j* (v*_i_* and v_j_) [25]. For example, in the grid-like biochemical reaction mesh network of Fig.1A, a direct path length from E636 and F649 was zero because there was a CH-π interaction between them. However, there was another shortest return path from F649 to Y627 and back to E636 via a π–π interaction. Because eight residues, ^628^NSLYSTCL^635^, between Y627 and E636 in the grid did not engage in non-covalent interactions, the grid size was 8. After each non-covalent interaction was tracked by a grid size and the uncommon sizes were marked in black, a grid with an x-residue or atom size was denoted as Grid_x_, and the total non-covalent interactions and grid sizes along the gating pathway of one subunit were calculated and shown in black and blue circles beside the mesh network map, respectively, for the calculation of the systematic thermal instability.

**Fig. 1.**
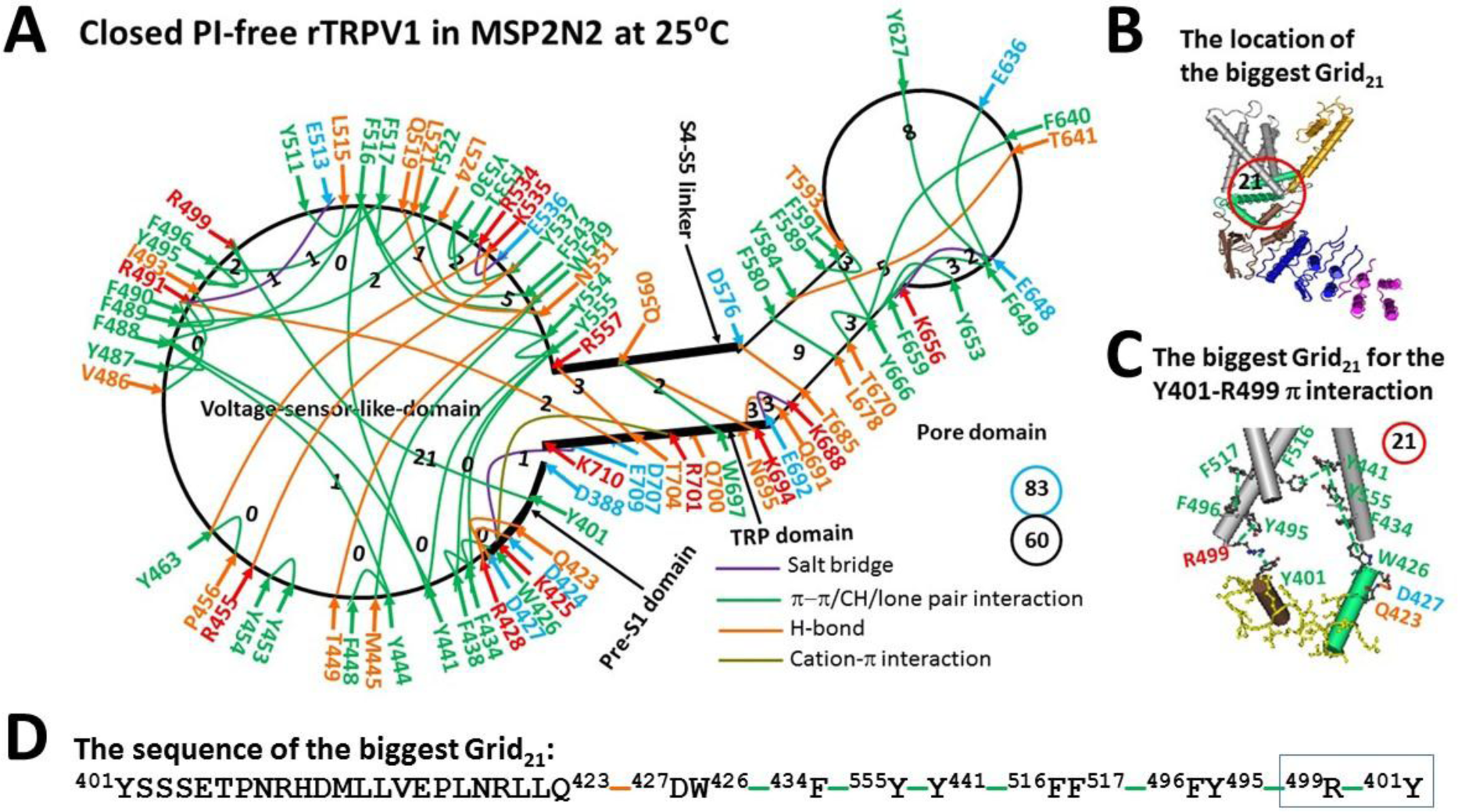
The grid-like non-covalently interacting mesh network along the gating pathway of PI-free rTRPV1 in the closed state at 25 °C. **A**, The topological grids in the systemic fluidic grid-like mesh network. The cryo-EM structure of one subunit in closed rTRPV1 with capsaicin bound at 25 °C (PDB ID, 7LPB) was used for the model [14]. The pore domain, the S4-S5 linker, the TRP domain, the VSLD and the pre-S1 domain are indicated in black. Salt bridges, π interactions, and H-bonds between pairing amino acid side chains along the gating pathway from D388 to K710 are marked in purple, green, and orange, respectively. The grid sizes required to control the relevant non-covalent interactions were calculated with graph theory and labeled in black. The total grid sizes and grid size-controlled non-covalent interactions along the gating pathway are shown in the blue and black circles, respectively. **B,** The location of the biggest Grid_21_ in the TRP/pre-S1/VSLD/TRP interfaces. **C,** The structure of the biggest Grid_21_ with a 21-residue size in the pre-S1/VSLD interface to control the the R499-Y401 π interaction. **D,** The sequence of the biggest Grid_21_ to control the R499-Y401 π interaction in the blue box.

### 2.4 Equations

The temperature threshold (T_th_) was calculated from a melting temperature T_m_ of the biggest grid along the gating pathway using the following equation as described previously [22]:

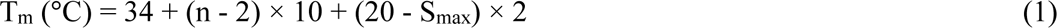

where, n is the total number of H-bonds equivalent to the grid size-controlled non-covalent interactions in the biggest grid, and S_max_ is the size of the biggest grid.

In either gating state, the systematic thermal instability (T_i_) along the gating pathway was defined using the following equation as described previously [22]:

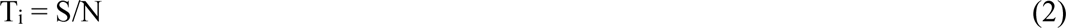

where, S is the total grid sizes along the gating pathway of one subunit in a given gating state; N is the total non-covalent interactions along the same gating pathway of one subunit in the same gating state.

For enthalpy-driven TRPV1 opening, if a thermosensitive TRPV1 channel changes from a fully closed state to a fully open state within a temperature range ΔT, and if the chemical potential of a grid is defined as the maximal potential for equivalent residues in the grid to form a tight β-hairpin with the smallest loop via non-covalent interactions [26], the grid-based structural thermo-sensitivity (Ω_ΔT_) of a single ion channel can be defined and calculated using the following equations:

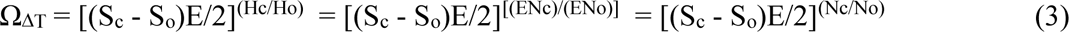

where, N_c_ and N_o_ are the total non-covalent interactions and H_c_ and H_o_ are the total enthalpy included in them in the closed and open states along the same gating pathway of one subunit, respectively; S_c_ and S_o_ are the total grid sizes along the same gating pathway of one subunit in the closed and open states, respectively. E is the energy intensity of a non-covalent interaction in a range of 0.5-3 kJ/mol. Usually, E is 1 kJ/mol [27]. Thus, Ω_ΔT_ factually reflects a heat-evoked change in the total chemical potential of grids upon a heat-evoked change in the total enthalpy included in non-covalent interactions from a closed state to an open state along the same gating pathway of one subunit. In the putatively inactivated state, Ω_ΔT_ was calculated when the parameters of the closed stated in this equation were replaced with those of the putatively inactivated state.

When ΔT=10 °C, Ω_10_ could be comparable to the functional thermo-sensitivity or the temperature coefficient (Q_10_) of a single ion channel. Q_10_ during thermal activation was calculated using the following equation:

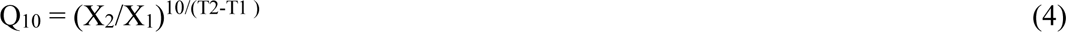

However, during thermal inactivation, it was calculated using the following equation:

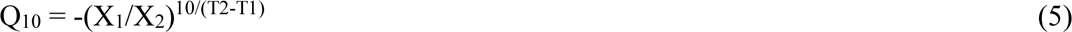

where, X_1_ and X_2_ are open probability (P_o_) values or reaction rates obtained at temperatures T1 and T2 (measured in kelvin), respectively.

## 3. RESULTS

### 3.1. Closed PI-free rTRPV1 at 25 °C Had the Biggest Grid_21_ for the calculated T_m_ 32 °C

The previous chimera studies between rTRPV1 and rTRPV2 demonstrated that the transmembrane segment 434-687 and the pre-S1 segment 401-415 may be involved in the regulation of the temperature threshold while the pre-S1 segment 358-434 plays a critical role in mediating the temperature sensitivity Q_10_. In contrast, an exchange of the C-terminal segment 688-838 has no effect [18]. On the other hand, the prior chimera investigations between heat-sensing TRPV1 and cold-sensing TRP melastatin 8 (TRPM8) indicated that the C-terminal including the TRP domain (693-710) is required for the polarity of thermal sensitivity [28]. Therefore, the segment from D388 in the pre-S1 domain to K710 in the TRP domain should be at least included as the gating pathway for the temperature threshold and sensitivity and the systematic thermal instability. At 25 °C, several kinds of non-covalent interactions between amino acid side chains along this gating pathway of PI-free rTRPV1 formed a biophysical network of topological grids in the closed state (**Fig.1A**).

First, H-bonds between different hydrophilic amino acid side chains contributed to the formation of network grids. In the pre-S1/VSLD interface, Q423-D427 and D424-R428 were paired for H-bonding. In the VSLD, T449-W549, R455-Y537-K535-Y463 and Q519-N551 were H-bonded in between. In the VSLD/S4-S5 linker/TRP/VSLD interfaces, the H-bonding pairs were R491-E709, R557-T704 and Q560-K694. In the pore domain and its interface with the TRP domain, pairs D576-T685, Y584-T641 and Q691-N695 were also engaged in H-bonds (**Fig. 1A**).

Second, aromatic π interactions also brought about network grids. Most of them were located in the VSLD and its interface with the pre-S1 domain. For example, Y401-R499-Y495, F434-W426/F438/Y555, F438-Y555, Y441-F488/F516/Y555, Y444-F448/F488, M445-W549, Y453-Y454, P456-Y463, V486-F490, Y487-F488/R491, F488-F516, F489-I493/L524, Y495/F517-F496, Y511-L515, F516-N551/Y554, F517-L521, F522-F543, Y530-F531/Y537 and Y554-Y555. In the S4-S5 linker/TRP interface, Q560 formed a CH-π interaction with W697. In the pre-S1/TRP interface, R701 produced a cation-π interaction with W426. In the pore domain, the π-interacting pairs had F580-L678, F589-T593, F591/T670-Y666, Y627/E636-F649, F640-Y666, and F649/Y653-F659 (**Fig. 1A**).

Third, salt-bridging pairs included K425-E709 in the pre-S1/TRP interface, R491-E513 and R534-E536 in the VSLD, E648-K656 in the outer pore, and K688-E692 in the S6/TRP interface (**Fig. 1A**).

Taken together, along the gating pathway from D388 to K710, sixty non-covalent interactions produced the total grid sizes as 83 (**Fig. 1A**). Thus, the systematic thermal instability (T_i_) of this closed PI-free-rTRPV1 was 1.38 (**Table 1**). Among the multiple grids, despite several smaller ones with 0- to 9-residues sizes, the biggest Grid_21_ was outstanding in the pre-S1/VSLD interface. It had a 21-residue size via the shortest path from Y401 to Q423, D427, W426, F434, Y555, Y441, F516, F517, F496, R499, and back to Y401 to control the R499-Y401 π interaction (**Figs. 1B-D**). Because two equivalent H-bonds sealed this biggest Grid_21_, the predicted melting temperature was about 32 °C (**Table 1**).

**Table 1.**
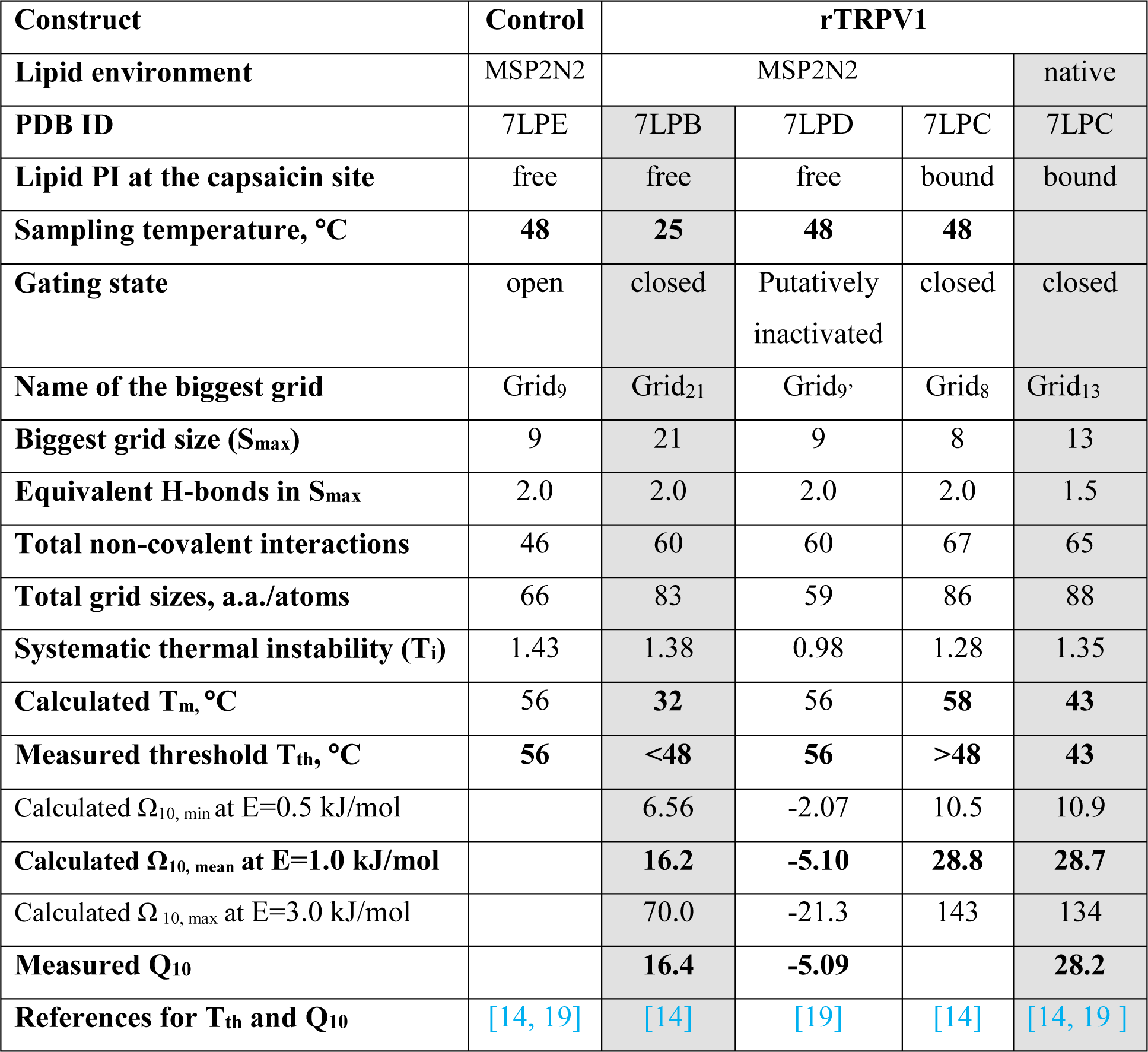
New parameters of the rTRPV1 bio-thermometer based on the grid thermodynamic model.

### 3.2. PI-free rTRPV1 Opened at 48 °C with the Melting of the Biggest Grid_21_

If the biggest Grid_21_ with a 21-residue size in rTRPV1 carries out the single-step melting reaction like DNA hairpins, the calculated melting temperature 32 °C should be enough to open it above 32 °C (**Table 1**). In agreement with this assumption, when PI-free rTRPV1 opened at 48 °C, the broken R499-Y401 π interaction resulted in the melting of the biggest Grid_21_ in the VSLD/pre-S1 interface together with several changes along the gating pathway.

In the pre-S1/VSLD/TRP interfaces, the K425-E709 salt bridge and the Q423-D427 and D424-R428 and R491-D707 H-bonds were disconnected. In the VSLD/S4-S5 linker/TRP interfaces, when the Y554-Y555 π interaction was disrupted, the H-bond moved from R557-T704 to Y554-T704 and the π interaction shifted from Y555-F438 to F559-F438. When the Q560-K694 H-bond was disrupted, the R557-E570 H-bond and the W697-R701 CH-π interaction were formed (**Fig. 2A**).

**Fig. 2.**
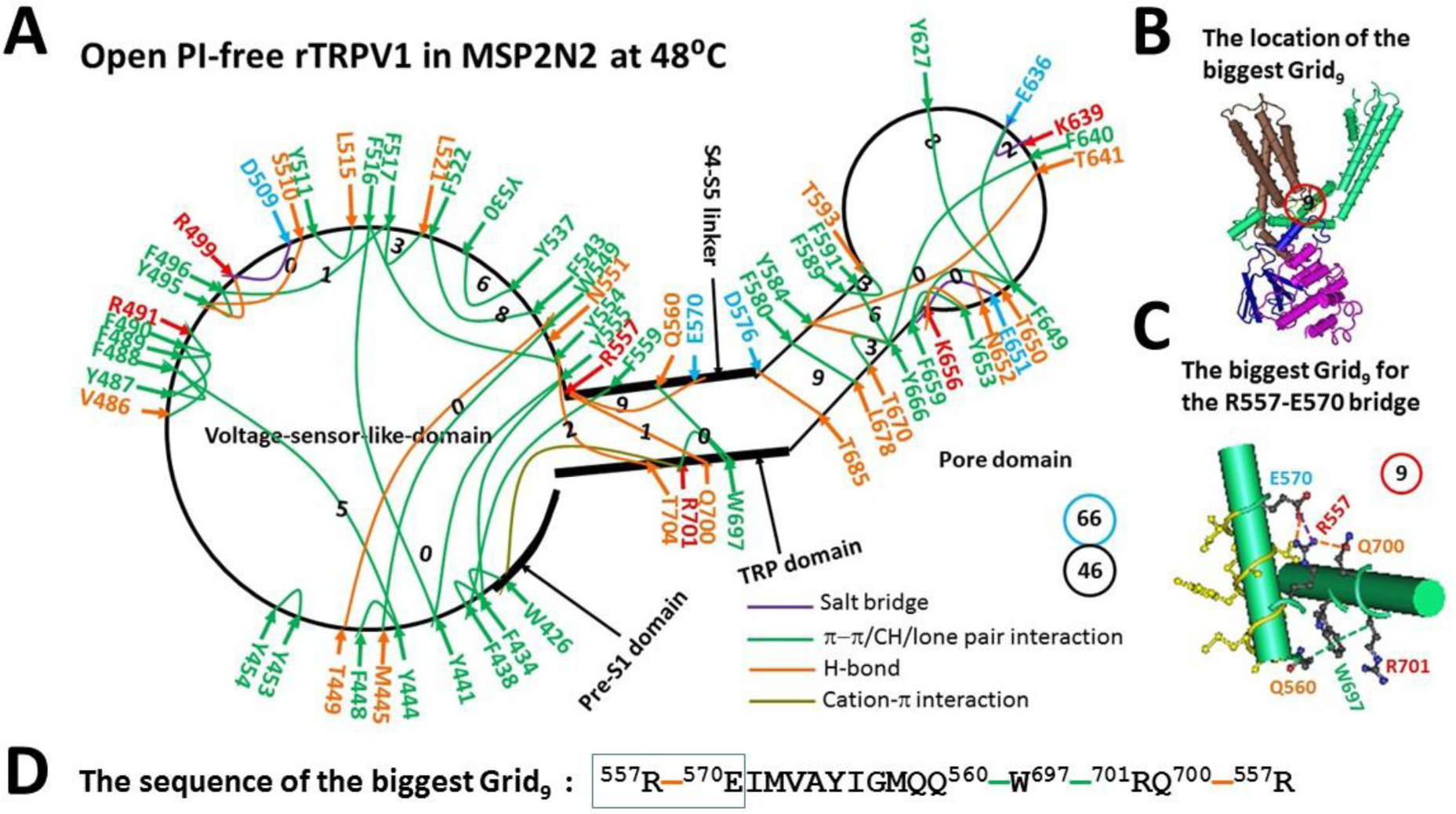
The grid-like non-covalently interacting mesh network along the gating pathway of PI-free rTRPV1 in the open state at 48 °C. **A**, The topological grids in the systemic fluidic grid-like mesh network. The cryo-EM structure of one subunit in open rTRPV1 with capsaicin bound at 48 °C (PDB ID, 7LPE) was used for the model [14]. The pore domain, the S4-S5 linker, the TRP domain, the VSLD and the pre-S1 domain are indicated in black. Salt bridges, π interactions, and H-bonds between pairing amino acid side chains along the gating pathway from D388 to K710 are marked in purple, green, and orange, respectively. The grid sizes required to control the relevant non-covalent interactions were calculated with graph theory and labeled in black. The total grid sizes and grid size-controlled non-covalent interactions along the gating pathway are shown in the blue and black circles, respectively. **B,** The location of the biggest Grid_9_. **C,** The structure of the biggest Grid_9_ with a 9-residue size in the VSLD/S4-S5 linker/TRP interfaces to control the stimulatory R557-E570 H-bond. **D,** The sequences of the biggest Grid_9_ with a 9-residue size to control the R557-E570 H-bond in the blue boxes, respectively.

When this conformational change extended to the VSLD, the R491-E513 and R534-E536 salt bridges, the R455-Y537-K535-Y463 and Q519-N551 H-bonds, and the Y441/F516-F488 and F489-I493 and Y530-F531 π interactions were disconnected. Meanwhile, the new Y495-R499/S510 H-bonds and the new R499-D509 salt bridge and the new F489-F490 π interaction emerged (**Fig. 2A**).

When this conformational change extended to the pore domain and its interfaces with the S4-S5 linker and the TRP domain, the E648-K656 and K688-E692 salt bridges and the Q691-N695 H-bond were disrupted. In the meanwhile, the new Y584-Y666 and T650-N652 and Y653-K656 H-bonds and the new E651-K656 salt bridge were born (**Fig. 2A**).

By all accounts, the total non-covalent interactions and grid sizes along the gating pathway from D388 to K710 dramatically decreased from 60 and 83 to 46 and 66, respectively (**Fig. 2A**). As a result, the systematic thermal instability (T_i_) of the gating pathway was increased from 1.38 to 1.43 along with the mean structural thermo-sensitivity Ω_10_ (16.2), which was close to the experimental Q_10_ (16.4) (**Table 1**) [14]. On the other hand, when two equivalent H-bonds sealed the biggest Grid_9_ with a 9-residue size in the VSLD/S4-S5 linker/TRP interfaces to control the stimulatory R557-E570 H-bond, the melting temperature was about 56 °C (**Fig. 2B-C, Table 1**).

### 3.3. PI-bound Native rTRPV1 Had the Increased Temperature Threshold and Sensitivity

When the resident PI occupies the best known vanilloid site, rTRPV1 in MSP2N2 cannot open even at 48 °C. This closed state of PI-bound rTRPV1 is almost identical between 4 °C and 48 °C [14]. However, once the PI lipid linked R409, D509, R557, E570 and Q700 together via H-bonds, there were several differences between PI-bound rTRPV1 at 48 °C and PI-free rTRPV1 at 25 °C (**Figs. 3A-B**).

**Fig. 3.**
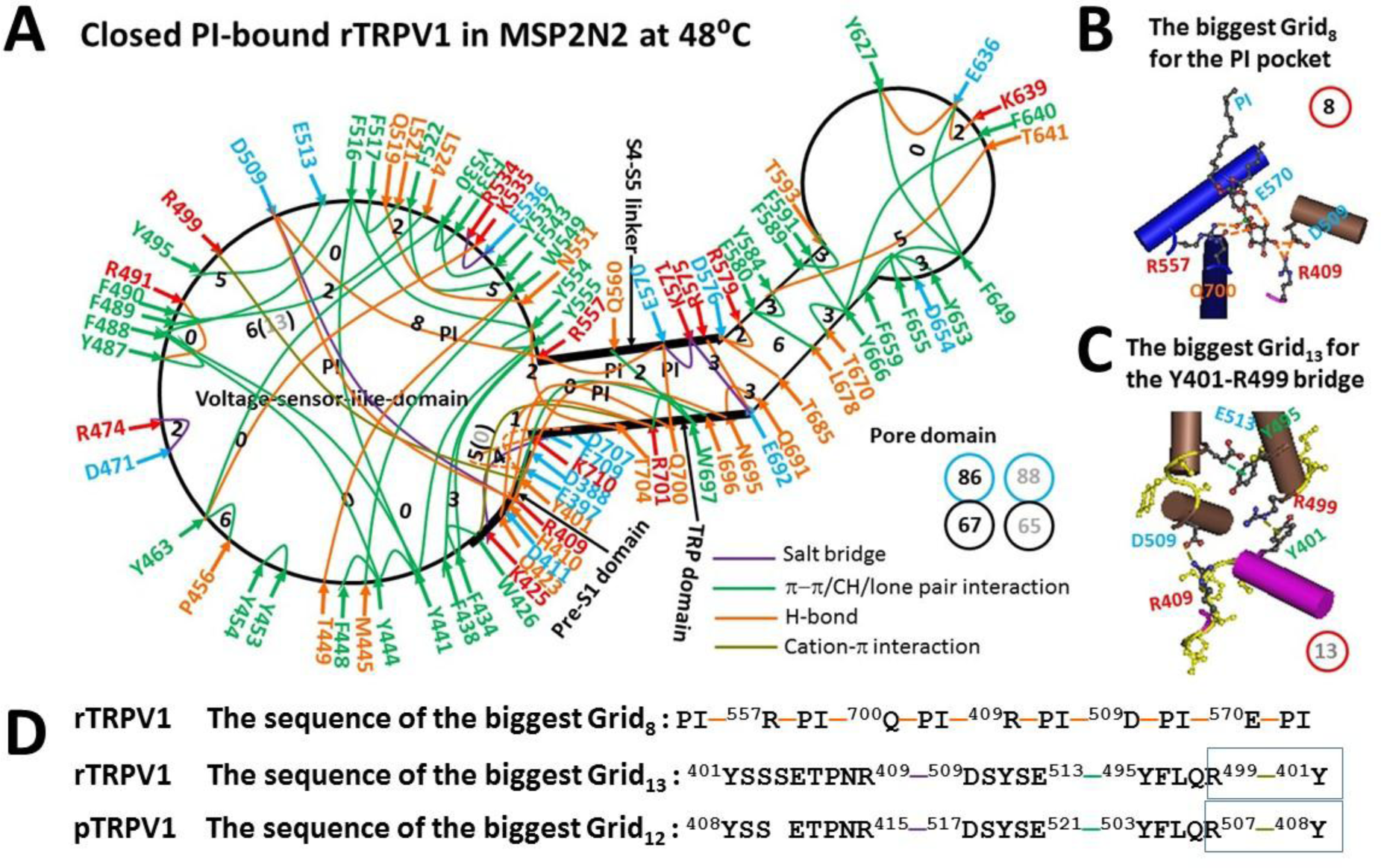
The grid-like non-covalently interacting mesh network along the gating pathway of PI-bound rTRPV1 in the closed state at 48 °C. **A**, The topological grids in the systemic fluidic grid-like mesh network. The cryo-EM structure of one subunit in closed rTRPV1 with PI bound at 48 °C (PDB ID, 7LPC) was used for the model [14]. The pore domain, the S4-S5 linker, the TRP domain, the VSLD and the pre-S1 domain are indicated in black. Salt bridges, π interactions, and H-bonds between pairing amino acid side chains along the gating pathway from D388 to K710 are marked in purple, green, and orange, respectively. The grid sizes required to control the relevant non-covalent interactions were calculated with graph theory and labeled in black. The total grid sizes and grid size-controlled non-covalent interactions along the gating pathway are shown in the blue and black circles, respectively. The dashed lines are absent in the native lipid system. The putative grid sizes in the native lipid system are labled in grey. **B,** The structure of the biggest Grid_8_ with a 8-atom size for the PI pocket. **C,** The structure of the biggest Grid_13_ with a 13-residue size in the VSLD/pre-S1 interface to control the Y401-R499 cation-π interaction near PI. **D,** The sequences of the biggest Grid_8_, Grid_13_, Grid_12_ with 8-carbon, 13-residue, and 12-residue sizes to control the PI pocket in rTRPV1, the Y401-R499 π interaction in rTRPV1, and the Y408-R507 π interaction in pTRPV1, respectively. The controlled bridges are marked in the blue rectangles.

In the pore domain and its interfaces with the S4-S5 linker and the TRP domain, the E648-K656 and K688-E692 salt bridges were disrupted with the formation of the D576-R579 and Y627-E636-K639 and E692-R575 H-bonds, the K571-E570/E692 salt bridges and the D654-F655 π interaction. In the VSLD, the R491-E513 salt bridge, the R455-Y537-K535 H-bonds, and the F434-F438 and V486-F490 and F489-I493 and Y495-F496/R499 and Y511-L515 π interactions were disrupted. In the meanwhile, the Y441-Y444 and Y463-Y530 and F489-F490 and Y495-E513 π interactions were formed with the Y487-R491 π interaction changing to an H-bond (**Fig. 3A**).

In the VSLD/TRP interface, when the H-bond shifted from R557-T704 to Y554-T704, the R491-D707 H-bond was disconnected. In the S4-S5 linker/TRP interface, the Q560-K694 H-bond was also broken but the W697-R701 CH-π interaction appeared. In the pre-S1/TRP interfaces, the new E397-K710, Y401-D707, D411-N695 and Q423-R701 H-bonds and the new H410-I696 π interaction were added. Finally, in the pre-S1/VSLD interface, the Q423-D427 and D424-R428 H-bonds were disrupted but a R409-D509 salt bridge was created (**Fig. 3A**). In this case, the total non-covalent interactions and grid sizes along the gating pathway were 67 and 86, respectively. Thus, the systematic thermal instability (T_i_) of PI-bound rTRPV1 in MSP2N2 decreased from 1.38 to 1.28 (**Table 1**). If the same open state was employed as a control, the mean structural thermo-sensitivity Ω_10_ would be about 28.8, which was comparable to the Q_10_ value 28.2 (**Table 1**) [14]. However, the tight interaction of rTRPV1 with MSP2N2 allowed the PI pocket to be the biggest Grid_8_ with an 8-atom size (**Fig. 3B**). When two equivalent H-bonds sealed this grid, the calculated melting temperature was about 58 °C (**Table 1**). That may be why the PI-bound rTRPV1 in MSP2N2 cannot open at 48 °C [14].

Once the E397-K710 and Y401-D707 H-bonds had been broken in a normal lipid environment, the biggest Grid_13_ with a 13-residue size via the shortest path from Y401 to R409, D509, E513, Y495 and R499 and back to Y401 would provide a melting temperature 43 °C as a normal threshold to disrupt the Y401-R499 π interaction and the related R409-D509 salt bridge so as to release the nearby PI lipid from the vanilloid site for rTRPV1 opening (**Table 1, Figs. 3A-D**). In that case, the total non-covalent interactions and grid sizes along the same gating pathway would be 65 and 88, respectively, so that the systematic thermal instability (T_i_) of PI-bound rTRPV1 in the native lipid environment would be 1.35, which was similar to that in PI-free rTRPV1 at 25 °C (**Table 1**). If the same open state had been employed as a control, the mean structural thermo-sensitivity Ω_10_ would be about 28.7, which was also comparable to the Q_10_ value 28.2 (**Table 1**) [14]. In that regard, the E397-K710 and Y401-D707 H-bonds may inactivate rTRPV1 from a closed state in the presence of MSP2N2.

### 3.4. PI-free rTRPV1 Inactivated Putatively at 48 °C after 30 s

In addition to an open state at 48 °C, a new state at 48 °C after 30 s for PI-free rTRPV1 (EMD 23478, PDB-7LPD) has been proposed as a pre-open closed state when the major class for 10 s (EMD 23477) was similar to the minor class for 30 s [14]. If the open state was used a control, the same broken Y401-R499 π interaction in Grid_21_ resulted in different changes along the gating pathway. For example, the Y401-E397 π interaction was present in the pre-S1 domain. On the other hand, in the VSLD, the R499-D509 salt bridge and the Y495-S510 H-bond, together with the F434-F438 and F489-F490 and Y495-F496 and F517-F496/L521 π interactions, were disrupted. Meanwhile, the F438-Y555 and Y441-F488 and P456-Y463 and F489-I493 and Y530-F531 and Y554-Y555 π interactions and the Y463-K535/Y537 H-bonds and the R534-E536 salt bridge were restored with the new Y441-Q519 and R491-Q494 H-bonds and the Y495-E513 π interaction. As a result, the F559-F438 π interaction in the S4-S5 linker/VSLD interface was broken, the H-bond moved from R557-E570 to E570-Q700 in the VSLD/S4-S5 linker/TRP interfaces, and the W697-R701 CH-π interaction became a strong cation-π interaction. Finally, when the conformational change extended to the outer pore, the E636-K639 salt bridge changed to an H-bond and the H-bonding pair moved from T650-N652 to E651-Y653. Of special note, the formation of the H410-E692 π interaction and the N695-H410 and Y401-D707 H-bond and the E397-K710 salt bridge in the pre-S1/TRP interface was a reminiscent of the similar or same non-covalent interactions in PI-bound inactivated rTRPV1 in the presence of MSP2N2 at 48 °C (**Figs. 3A, 4A**).

**Fig. 4.**
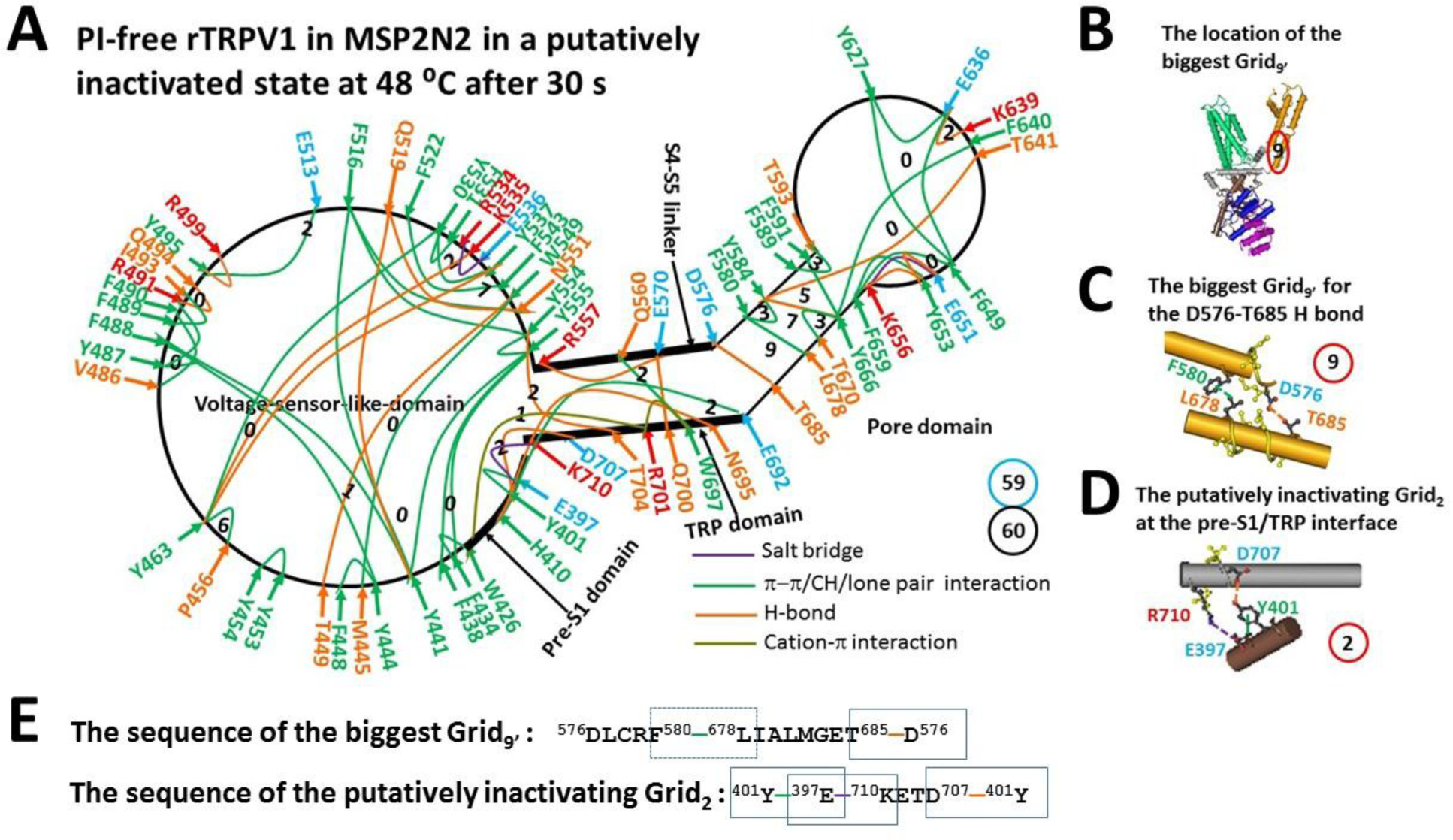
The grid-like non-covalently interacting mesh network along the gating pathway of PI-free rTRPV1 in a putatively inactivated state at 48 °C. **A**, The topologocal grids in the systemic fluidic grid-like mesh network. The cryo-EM structure of one subunit of rTRPV1 with capsaicin bound at 48 °C for 30 s (EMD 23478; PDB ID, 7LPD) was used as a model for a putatively inactivated state [14]. The pore domain, the S4-S5 linker, the TRP domain, the VSLD and the pre-S1 domain are indicated in black. Salt bridges, π interactions, and H-bonds between pairing amino acid side chains along the gating pathway from D388 to K710 are marked in purple, green, and orange, respectively. The grid sizes required to control the relevant non-covalent interactions were calculated with graph theory and labeled in black. The total grid sizes and grid size-controlled non-covalent interactions along the gating pathway are shown in the blue and black circles, respectively. **B,** The location of the biggest Grid_9‘_. **C,** The structure of the biggest Grid_9‘_ in the S5/S6 interface to control the D576-T680 H-bond and the F580-L678 π interaction. D, The putatively inactivating Grid_2_ to govern the Y401-D707 H-bond and the E397-K710 salt bridge. E, The sequences of the biggest Grid_9‘_ and the putatively inactivating Grid_2_ with 9- and 2-residue sizes to control the non-covalent interactions in the blue rectangles, respectively.

Taken together, when the biggest Grid_9’_ in the pore domain had a 9-residue size to control the D576-T685 H-bond and the F580-L678 π interaction at the S5/S6 interface via the shortest path from D576 to F580, L678, T685 and back to D576 (**Figs. 4A-C, E**), the resultant melting temperature T_m_ was about 56 °C, the same as that of the open state (**Table 1**). On the other hand, when the total non-covalent interactions and grid sizes along the gating pathway from D388 to K710 significantly changed from 46 and 66 to 60 and 59, respectively (**Fig. 2A, 4A**), the calculated Ω_10_ had a range from −2.07 to −21.3 and a mean value −5.10 (**Table 1**), which was close to the experimental Q_10_ of the inactivated state (−5.09). [19] Further, this new state had the lower systematic thermal instability (T_i_ = 0.98) than those of the closed state at 25 °C and the open state at 48 °C (Ti=1.38, 1.43, respectively) (**Table 1**). Finally, the putatively inactivating Y401-D707 and E397-K710 bridges as found in PI-bound rTRPV1 at 48 °C were also present in this new state (**Figs. 1A, 3A, 4D-E**). Accordingly, the major class for 10 s may be a pre-open closed state but the similar minor class for 30 s may serve as a putatively inactivated state from the pre-open closed state and would become a major class after 50 s (**Fig. 5A**). [14, 20]

**Fig. 5.**
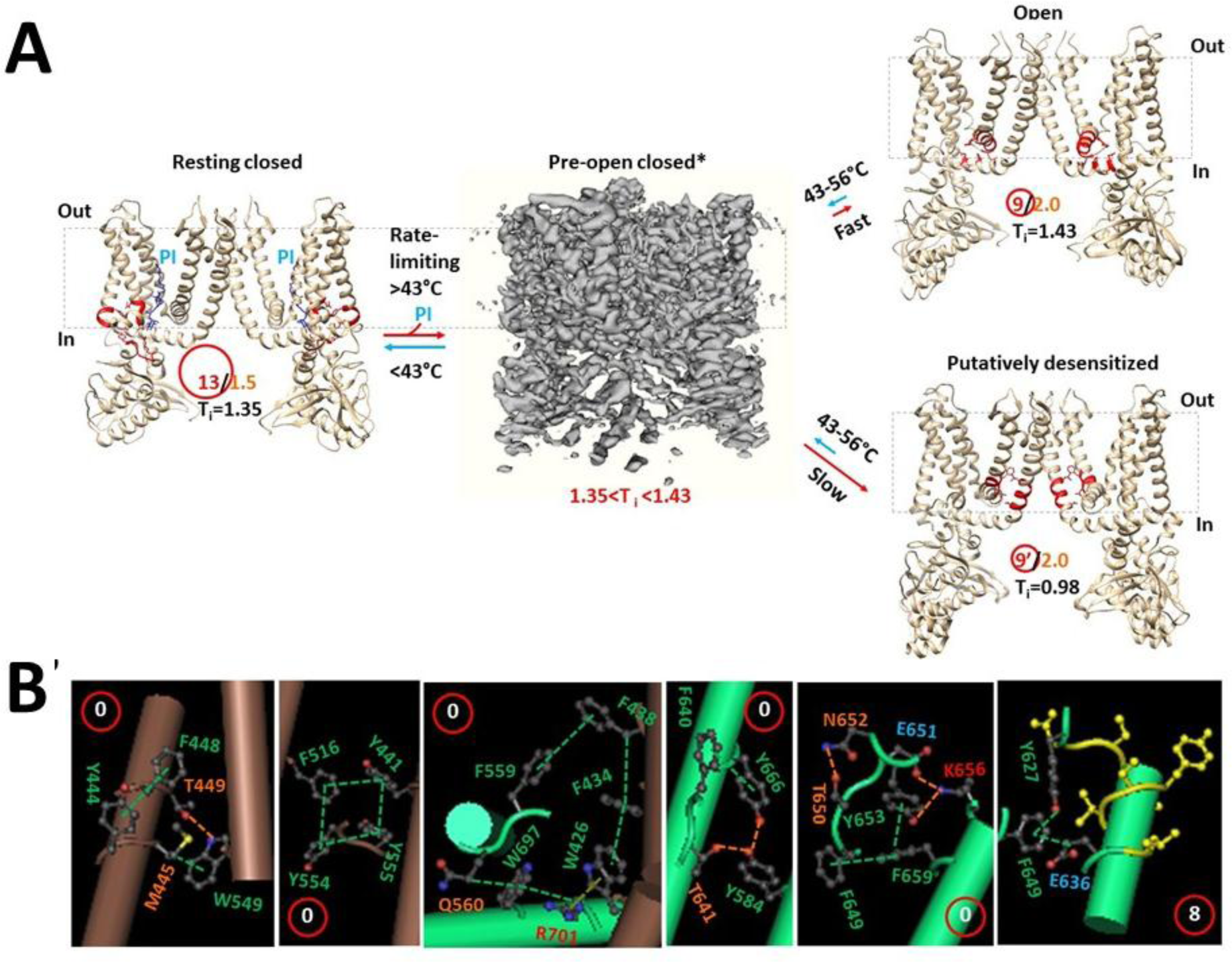
The tentative model for TRPV1 opening and desensitization upon the heat-evoked melting of the biggest grid. **A.** The homo-tetrameric cryo-EM structures of rTRPV1 in the resting closed, open, putatively-inactivated and pre-open closed states (PDB ID:7LP9, 7LPE, 7LPD and EMD-23477, respectively) were used for the model [14]. For a convenient view, only two opposite subunits are shown except for the tetrameric pre-open closed state. The dashed rectangles are the membrane areas. In the presence of the PI lipid (blue) at the capsaicin site, rTRPV1 has 1.5 equivalent H-bonds (orrange) to seal the biggest Grid_13_ (red) with a 13-residue size (red) in the VSLD/pre-S1 interface so that rTRPV1 is closed below a threshold 43 **°**C. When the temperature increases above the threshold 43 **°**C, rTRPV1 swiftly opens from a pre-open closed state within 30 s upon the melting of the rate-limiting biggest Grid_13_ in the VSLD/pre-S1 interface along with the release of the PI lipid from the vanilloid site [14, 20]]. In the meantime, the biggest Grid_9_ (red) with a 9-residue size (red) and two equivalent H-bonds (orrange) is formed in the VSLD/S4-S5 linker/TRP/VSLD interfaces in favor of the open state below a T_m_ 56 **°**C. On the other hand, the channel may also slowly inactivate from the same pre-open closed state after 30 s [14, 20]. Clearly, the mean systematic thermal instability (T_i_) between 1.35 and 1.43 of the pre-open closed state may faciliate the fast opening. However, the lower T_i_ (0.98) of the putatively inactivate state may allow rTRPV1 to bypass the open state and to inactivate slowly from the pre-open closed state in a irreversable manner at prolonged temperature between 43 °C and 56 **°**C [19, 20]. **B,** The proposed smaller grids for TRPV1 gating and heat efficacy. The grid sizes are shown in the red circles.

## 4. DISCUSSION

TRPV1 is designed and created as a bio-thermometer in the thermostable system to detect noxious heat and to protect human or animal bodies from heat damage [1-4]. To this end, it should meet several protein engineering requirements. First, it should be an integral membrane protein with a relative high physicochemical stability or heat capacity to avoid thermal denaturation. Second, it should be an ion channel that can directly couple a temperature signal to an electronical signal reporter for a readout by a primary afferent sensory neuron. Third, its opening from the closed state within 10 °C apart should have a high Q_10_ with a steep response to a small change in a surrounding predetermined temperature. Fourth, it should have a heat switch with two temperature threshold limits for two gating states, one for activation to sense noxious heat and the other for inactivation to alleviate pain by exploiting the use-dependent desensitization upon constant or repeated noxious heat stimuli. In the biochemical and biophysical world, a DNA hairpin-like topological thermal grid in a systemic fluidic grid-like non-covalently interacting mesh network may be an ideal tool for TRPV1 to achieve them. The grid thermodynamic model, once has been successfully used to pinpoint the structural motifs as the local thermal sensors for the changes in the structural thermo-stability and the functional thermo-activity of a biological macromolecule aldolase B [22], could be applied to this computational study to uncover the roles of the biggest and smaller grids in heat-evoked activation, high temperature sensitivity Q_10,_ and use-dependent desensitization.

### 4.1 The Role of the Biggest Grids in Heat-sensing

This computational study has identified five biggest grids in rTRPV1. Like a DNA hairpin or aldolase B [21-22], at a given physiological salt concentration (150 mM NaCl), when non-covalent interactions made the biggest grid along the gating pathway from D388 in the pre-S1 domain to K710 in the TRP domain of rTRPV1, the calculated melting temperature threshold (T_m_) values were exactly close to the measured T_th_ values (Table 1). Therefore, a single-step melting reaction of the biggest grid may be rate-limiting for the specific temperature-dependent threshold (T_th_) in rTRPV1. In other words, those biggest grids may serve as the specific local thermal sensors for different gating states. Grid_21_ and Grid_13_ may be allosterically coupled to the lower S6 gate via the R499-Y401 π interaction in the VSLD/pre-S1 interface. Therefore, their melting disrupted this non-covalent bridge, allowing PI-free and PI-bound rTRPV1 to open above 32 °C and 43 °C, respectively (**Table 1, Figs. 1-3**). In agreement with this proposal, in the presence of the putative biggest Grid_12_ with a 12-residue size via the shortest path from Y408 to R415, D517, E521, Y503, R507 and back to Y408 (**Fig. 4D**), platypus (*Ornithorhynchus anatinus*) TRPV1 (pTRPV1) decreases an activation threshold by about 3 °C [20]. When the salt concentration was lowered down from 150 mM to 130 mM, the threshold may also decrease from 42 °C to 37.7 °C [8, 20]. Furthermore, the turret deletion or replacement with glycine between G603 and N626, or the intracellular niacin or extracellular acidic pH 6.3 or the E600K mutation decreases the temperature threshold of mouse TRPV1 (mTRPV1) or rTRPV1 to a different extent [8, 22, 29-31], possibly by weakening the corresponding R500-Y402 π interaction in the VSLD/pre-S1 interface, or releasing the PI from the vanilloid site [16], or generating another new biggest grid with a size larger than 13 residues along the gating pathway from D389 to K711. Of special note, the N-terminal S124N/Q188E mutation may change the thermal sensor via the pre-S1/TRP interface and thus lose the temperature sensitivity [13, 32].

Following the melting of the biggest Grid_21_ or Grid_13_, Grid_9_ may favor the open state by uplifting the S4-S5 linker against the S6 gate via the stimulatory R557-E570 H-bond in the VSLD/S4-S5 linker interface. As the mutation R557E/A/L or E570L/A can disrupt the stimulatory R557-E570 H-bond, it is reasonable that the R557E or E570L mutant leads to a complete loss of function, or mutants R557A/L and E570A or nearby M572A have low heat sensitivity, or E570A increases the threshold from 41.5 °C to 44.3 °C [14, 33, 34]. Since Grid_9_ had an upper melting temperature limit 56 °C to keep the open state of rTRPV1 (**Figs. 2, 5A, Table 1**), it is reasonable that rTRPV1 exhibits the maximal activity at about 56 °C in response to a temperature ramp from 20 °C to 60 °C [19].

Unlike Grid_9_, Grid_9’_ was present in all the states of PI-free rTRPV1 unless the D576-R579 H-bond in the closed PI-bound rTRPV1 reduced the size to 6 residues (**Figs. 1A, 2A, 3A, 4A**). Since the residues in this grid are highly conserved in TRPV1-6, it may serve as a key common anchor to maintain an active pore domain [11, 35]. Any thermostable perturbation in this anchor grid may induce partial thermal inactivation of the ion conduction pathway from an open state, leading to flickering opening [22, 23]. In direct line with this notion, the mutant R579E is not functional and the D576N/R or R579A mutant is less sensitive to heat than wild type rTRPV1 [33]. In addition, the loss of heat sensitivity was also found in the T680A mutant of mTRPV3 (T685 in rTRPV1) [36].

Taking all these into account, rTRPV1 may have a molecular thermometer range from 43 °C to 56 °C to monitor noxious heat (**Fig. 5A**). Further structural and electrophysiological experiments are necessary to examine the roles of the Y401-R499 π interaction and the R409-D509 salt bridge in securing the normal temperature threshold 43 °C for rTRPV1 opening.

### 4.2 The Role of Smaller Grids in Stabilizing Heat Efficacy

In addition to the above biggest grids for heat-sensing, several smallest grids with a zero-residue size may also be required for channel gating. The first via the shortest path from Y444 to M445, W549, T449, F448 and back to Y544 was highly conserved in the VSLD of all the gating states and thus may serve as a static anchor (**Figs. 1-5**). The second via the shortest path from Y441 and through F516 and Y554 and Y555 and back to Y441 in the VSLD was shared by both closed and open states (**Figs 1-2, 5B**), and the related aromatic residues were highly conserved in thermosensitive TRPV1-4 but not in non-thermosensitive TRPV5-6 [11, 35]. Such tight π interactions may allow both smallest grids to serve as relatively stationary heat fuses or anchors against which the S4–S5 linker can uplift in favor of TRPV1 opening. In support of this notion, the Y441S, Y444S, M445W, Y554A or Y555S mutation leaves the channel not functional [33, 37]. The third linked W426 on pre-S1, F434 and F438 on S1, F559 and Q560 in the S4-S5 linker, and W697 and R701 in the TRP domain together via the shortest path. This Grid_0_ may play a critical role in stabilizing the open state (**Figs. 2, 5B**). The fourth via the shortest path from Y584 to T641 and F640 and Y666 and back to Y584 was present in the pore domain of the open and putatively desensitized states. (**Fig. 5B**). The residues for this smallest Grid_0_ were also highly conserved in TRPV1-4 but not in TRPV5-6 [11, 35]. In agreement with its importance, the Y666A mutation is not functional while the F640L or T641S mutation decreases the temperature threshold to allow either to be constitutively active at room temperature [38-39]. The fifth may stabilize the open state by linking F649, F659, Y65, K656, E651, N652, T650 together against a highly-conserved anchor Grid_8_ via the shortest path from Y627 to F649 and E636 and back to Y627 in the pore domain (**Figs. 2A, 5B**). It has been reported that the N628K or N652T or Y653T or combined mutation increases the temperature threshold, and the triple mutant N628K/N652T/Y653T has the longer opening durations. [7] Furthermore, the N628L mutation greatly decreases the heat sensitivity (Q_10_ ∼ 2.6) but the N628D mutation has the opposite effect (Q_10_ ∼ 86) [14]. possibly by mediating the thermal stability of these two smaller grids through ionic interactions with nearby E636. Notably, although the deletion of the turret (Δ604-626) still allows the mTRPV1 mutant construct to be activated by heat, the binding of external double-knot toxin or vanilla-toxin (DkTX) to the outer pore prevents heat activation [9, 15], possibly by affecting the grids in the outer pore (**Fig. 5B**).

Taken as a whole, while the biggest grids could serve as local thermal sensors to determine the initial temperature thresholds for gating transitions, smaller ones may play a critical role in stabilizing the final heat efficacy. Therefore, all the grids from the biggest to the smallest along the activation pathway from D388 to K710 may be needed for the high temperature sensitivity.

### 4.3 The Structural Origin of the Temperature Sensitivity

When PI-free rTRPV1 opens at 48 °C from a closed state at 25 °C, the opening of multiple grids may increase entropy but decrease enthalpy as a result of the decrease in the total non-covalent interactions from 60 to 46 along the gating pathway from D388 to K710 (**Figs. 1-2**). Thus, a small decrease in the Gibbs free energy may be enough for intra-molecular gating rearrangements in the grid-like non-covalent interaction mesh networks to activate TRPV1. However, even if multiple short links in grids were included, such a small decrease may be insufficient to produce a high Q_10_ for noxious heat detection. Therefore, a heat-induced decrease in the total chemical potential of all the grids upon the heat-evoked decrease in the total enthalpy included in the non-covalent interactions along the same gating pathway from a closed state to an open state should be considered. In consonance with this proposal, when the total grid sizes dramatically changed along with the total non-covalent interactions, the resultant mean structural thermo-sensitivity (Ω_10_) was comparable to the functional thermo-sensitivity (Q_10_). For PI-free rTRPV1, the calculated mean Ω_10_ was about 16.2, while the experimental Q_10_ is about 16.4. For PI-bound rTRPV1, the predicted mean Ω_10_ was about 28.7 which was close to the experimental Q_10_ is 28.2 [1, 14]. When the intensity of a non-covalent interaction ranged from the minimal 0.5 kJ/mol to the maximal 3 kJ/mol, the normal Ω_10_ for channel opening could be in a range from 10.9-134 (**Table 1**). Therefore, any measured Q_10_ values for native rTRPV1 or mTRPV1 may be reasonable in this range [1, 8, 14, 18, 20].

It should be noteworthy that even if heat-evoked TRPV1 opening is enthalpy-drive, an increase in heat capacity (love) or energy density as a result of the decrease in the total grid sizes from a closed state to an open state along the same gating pathway may be still necessary to secure a minimal decrease in the Gibbs free energy for channel opening with a maximal Q_10_ value [40].

### 4.4 The Structural Motif for Use-dependent Desensitization

TRPV1 is characterized by intriguing use-dependent desensitization. However, the underlying structural motif is missing. In this computational study, the similar experimental T_th_ (>41 °C) for rTRPV1 activation and inactivation, [18-19] together with the similar structures between the pre-open closed state and the putatively inactivated state, [14] and the same calculated melting temperature 56 °C of the biggest grids in both putatively inactivated and open states (**Table 1**), suggests that, like the inactivation from a pre-open state of the Kv4 channel complexes [41], those two gating states may originate from the same pre-open closed state upon the release of PI from the vanilloid site (**Fig. 5A**). However, similar Ω_10_ (−5.10) and Q_10_ (−5.09) for inactivation was weaker than similar Ω_10_ (28.7) and Q_10_ (28.2) for channel opening (**Table 1**) [18]. Thereafter, channel opening may be faster than the putative inactivation (**Fig. 5A**). That may account for why the major class appears at 48 °C for 10 s but the similar minor class was present at 48 °C for 30 s and would become a major class after 50 s [14, 20].

On the other hand, among different gating states, the normal systematic thermal instability (T_i_) along the same gating pathway from D388 in the pre-S1 domain to K710 in the TRP domain was about 1.35 to 1.38 in the closed state no matter whether the PI was bound at the vanilloid site (**Table 1**). However, this value was significantly increased up to 1.43 in the open state but decreased to 0.98 in the putatively inactivated state (Table 1). Therefore, the pre-open closed state may have a T_i_ value between 1.35 and 1.43 in favor of fast activation but slow inactivation in an irreversible manner (**Fig. 5A**) [1, 8, 19-20].

It has been reported that the thermosensitive dynamic interaction between the N- and C-terminal domains (segments 1-433 and 688-839, respectively) of mTRPV1 but not pTRPV1 triggers a conformational rearrangement in the pore causing desensitization [20]. This computational study further indicated that the smaller Grid_2_ via the shortest path from Y401 to E397, K710, D707 and back to Y401 to control the E397-K710 or Y401-D707 H-bridge may have a very high melting temperature threshold (> 70 °C) to lock rTRPV1 in a putatively irreversible inactivated state below 70 °C at pH 7 (Figs. 4A, D, E), alleviating pain upon constant or repeated noxious heat stimuli-induced desensitization (**Table 1, Figs. 2-4**) [1, 8, 14, 19-20]. In accordance with this proposal, once PI-bound rTRPV1 was reconstituted in MSP2N2, the presence of those two bridges allowed the PI pocket at the vanilloid site to be the biggest Grid_8_ with the high melting temperature 58 °C to inhibit the release of PI from the vanilloid site at 48 °C for channel opening (**Table 1, Fig. 3**). In contrast, resiniferatoxin irreversibly opens TRPV1 without both desensitization and the Y401-D707 and E397-K710 H-bonds even at 48 °C [17, 42-43]. Furthermore, rTRPV1 without either or both of the Y401-D707 and E397-K710 H-bonds in three conformers at pH 6 still exhibits heat desensitization [29, 31]. However, in the Ca^2+^-free bath solution, when Y401-D707 and E397-K710 H-bonds are broken (PDB ID, 7L2O) at PH 5.5 [31], protons evokes nondesensitizing responses [29]. Taken together, the absence of a serine residue between S410 and E411 may not allow pTRPV1 to have the E404-K718 or Y408-D715 H-bond in the pre-S1/TRP interface to inactivate pTRPV1 (**Figs. 3E, 4D**) [20]. In other words, the deletion of S404 or the double E397A/Y401A mutation may stop the putative heat inactivation of rTRPV1, and the cryo-EM structure of pTRPV1 at pH 7 may have unique closed and open states but not an inactivated state around the specific heat threshold in the presence of 150 mM NaCl.

### 4.4 The Limitations of This Graphical Method

Although this graphical method has successfully predicted the temperature thresholds and sensitivities for the thermal activation and use-dependent inactivation of rTRPV1 or mTRPV1 and thus uncovered the related structural bioinformatics or digit codes, it may have some limitations. First, a high-resolution 3D structure is necessary to identify the systemic fluidic grid-like non-covalent interaction mesh network. However, such a strict requirement may not always be met for some membrane proteins including some TRP channels. Second, a suitable lipid system is needed to allow the structure of membrane protein to be captured in a correct functional state. Nonetheless, it is not easy to find an ideal lipid system to achieve it for structural biologists. For example, PI-bound rTRPV1 could not be opened in the presence of MSP2N2 even above the normal threshold 43 °C [1, 14]. Third, not all the biological macromolecules including proteins can have their accurate tertiary structures snapshoted at more than one temperatures from the lower to the higher. When the predicted T_m_ to stabilize the open or inactivated states was as high as 56 °C for rTRPV1 (Table 1), it may be a challenge to confirm it structurally.

## 5. CONCLUSION

The topological grids of a grid-like non-covalent interaction mesh network in an ion channel can be used to design a thermosensitive bio-thermometer with a high Q_10_ value. In addition to the ionic strength in the physiological range, the size of the biggest grid and the number and the intensity of the grid size-controlled non-covalent interactions between amino acid side chains in this grid can be used to tune the grid strength for different melting temperature thresholds to detect distinct minor environmental temperature changes. These non-covalent interactions include hydrophilic salt bridges and H-bonds as well as hydrophobic aromatic side chain π interactions. Both the biggest and smaller grids along the gating pathway are necessary to control channel gating transitions and the related thermal efficacy, respectively. Their opening and closure should be reversible and fast enough to have a very large temperature-dependent effect on gating switches, which can be easily coupled to an electrical readout from the primary afferent sensor neuron. What is more, the use-dependent desensitization may use gating state-dependent systematic thermal instability as a protective measure against noxious heat damage. In this regard, this computational study may render a broad structural basis for thermosensitive TRP channels to work as a molecular thermometer.

On the other hand, although a given non-covalent interaction’s spatial and geometric requirements have been well-understood, it is still hardly predictable in a complex grid-like temperature-dependent biochemical reaction mesh network. In that regard, this grid thermodynamic model, once examined by the temperature-dependent structural data of aldolase B and TRPV1, may provide more creative metrics for identifying causal structure– activity/selectivity relationships in a supramolecular architecture at a variety of ambient temperatures.

More importantly and significantly, this computational study demonstrated that the strength of a non-covalent interaction in a biological macromolecule can be regulated by the size of the grid sealed by that non-covalent interaction for the thermal adaption. This is an amazing finding regarding the origin of life at ambient temperature.

## Conventions and Abbreviations

cryo-EM: cryo-electron microscopy
DkTx: double-knot toxin or vanillotoxin
*k_d_*: the dissociation rate
Ω_10_: structural thermosensitivity
Q_10_: functional thermo-sensitivity
PI: phosphatidylinositol
P_o_: open probability
RTX, T_i_: thermal instability
T_m_: melting temperature
T_th_: temperature threshold
TRP: transient receptor potential
TRPM8: TRP melastatin 8
TRPV1: TRP vanilloid-1
TRPVi: (i=1; 2, 3, 4, 5, 6)
hTRPV1: human TRPV1
mTRPV1: mouse TRPV1
pTRPV1: platypus TRPV1
rTRPV1: rat TRPV1
VSLD: voltage-sensor-like domain

## Acknowledgements

The author’s own studies cited in this article were supported by NIDDK Grant (DK45880 to D.C.D.) and Cystic Fibrosis Foundation grant (DAWSON0210) and NIDDK grant (2R56DK056796-10) and American Heart Association (AHA) Grant (10SDG4120011 to GW).

## Author contributions

G. W. wrote the main manuscript text, prepared table 1 and figures 1-5, and reviewed the whole manuscript.

## Data availability statement

All data generated or analysed during this study are included in this published article.

